# Gigantic Genomes of Salamanders Indicate Body Temperature, not Genome Size, is the Driver of Global Methylation and 5-Methylcytosine Deamination in Vertebrates

**DOI:** 10.1101/2021.09.27.462072

**Authors:** Alexander Nichols Adams, Robert Daniel Denton, Rachel Lockridge Mueller

## Abstract

Methylation of cytosines at CpG dinucleotide sites silences transposable elements (TEs), sequences that replicate and move throughout genomes. TE abundance drives differences in genome size, but TE silencing variation across genomes of different sizes remains largely unexplored. Salamanders include most of the largest C-values — 9 to 120 Gb. We measured CpG methylation levels in salamanders with genomes ranging from 2N = ~58 Gb to 4N = ~116 Gb. We compared these levels to results from endo- and ectothermic vertebrates with more typical genomes. Salamander methylation levels are ~90%, higher than all endotherms. However, salamander methylation does not differ from the other ectotherms, despite a ~100-fold difference in nuclear DNA content. Because methylation affects the nucleotide compositional landscape through 5-methylcytosine deamination to thymine, we quantified salamander CpG dinucleotide levels and compared them to other vertebrates. Salamanders have comparable CpG levels to other ectotherms, and ectotherm levels are higher than endotherms. These data show no shift in global methylation at the base of salamanders, despite a dramatic increase in TE load and genome size. This result is reconcilable with previous studies by considering endothermy and ectothermy, which may be more important drivers of methylation in vertebrates than genome size.

## Introduction

Genomes are composed of sequences with radically different effects on organismal survival and reproduction. Essential genes are required to sustain life, and many are constitutively transcribed in all tissues. In contrast, transposable elements (TEs) — sequences capable of proliferation and movement throughout genomes — are mutagenic, and their transcriptional silencing is required to maintain germline integrity and reproduction (Bourque et al. 2018). Transcription or silencing of different DNA sequences is facilitated by conformational changes to chromatin, which is achieved through epigenetic modifications to both DNA and histones (Venkatesh and Workman 2015). The suppression of TEs relies on transcriptional silencing through methylation of cytosines (5mC) that are adjacent to guanines (i.e. CpG dinucleotide sites) (Bird 2002; Law and Jacobsen 2010; Fedoroff 2012; Venkatesh and Workman 2015; Deniz et al. 2019). TEs are a major determinant of overall genome size, which varies > 65,000-fold across eukaryotes (Pellicer et al. 2010). How TE suppression via methylation at CpG sites scales with increased TE load and genome size remains incompletely understood (Wolffe 1998; Jones and Wolffe 1999).

There have been few attempts to explore global methylation at the upper limits of genome size. In *Picea abies*, the Norway Spruce — which has an unusally large diploid genome size of 20 Gb — the percentage of methylated CpG sites is higher than that found in other monocots and eudicots with smaller genome sizes. CHG sites (where H is any nucleotide that is not a guanine) show the same pattern, whereas methylated CHH sites do not (Ausin et al. 2016). 5mC at CpG and CHG are both associated with TE silencing in plants, while 5mC at CHH sites is not, instead forming a barrier between heterochromatin and other genes (Kenchanmane Raju et al. 2019). This single data point suggests that, as genome size increases, the percentage of methylated cytosines at TE-silencing sites increases. However, to our knowledge, the relationship between genomic gigantism and global methylation in animals remains untested, as does the relationship in any taxon with a genome size above 20 Gb.

Global methylation levels can also affect the nucleotide compositional landscape because methylation causes an increase in the rate of specific mutations. Methylated cytosines at CpG sites are predisposed to undergo transition mutations from C to T via deamination (Ehrlich and Wang 1981). Recently, Zhou et al. (2020) demonstrated that the proportion of CpG sites decreases as genome size increases across animal species with genome sizes ranging from 89 Mb to 4 Gb, which was interpreted to reflect deamination-driven transition mutations occurring at higher frequency as TE loads and associated methylation levels increase. Whether this relationship holds for animals with gigantic genome sizes remains unknown.

The salamander clade is an appropriate model system for studying the epigenetic factors that control — or fail to control — TE activity in gigantic genomes because it includes diploid genomes that range from 9 to 120 Gb (Šímová and Herben 2012; Sun et al. 2012; Sclavi and Herrick 2019; Gregory 2020; Decena-Segarra et al. 2020). These gigantic and variably-sized genomes reflect the accumulation of extreme levels of TEs, associated with a shift in the dynamics of genome size evolution at the base of the clade ~200 mya (Batistoni et al. 1995; Sun and Mueller 2014; Liedtke et al. 2018). Additionally, there are several polyploid salamander species, providing a different mechanistic path to high levels of nuclear DNA by combining large genomes with increased chromosome copy number (Bogart et al. 2007, 2009).

Here, we quantify methylation levels across salamanders that encompass their upper range of genome sizes, and we compare these values to those of other vertebrates with more typically sized genomes. We include both endotherms (i.e. birds and mammals) and ectotherms (i.e. fish, other amphibians, reptiles) because body temperature is associated with differences in global methylation (Jabbari et al. 1997). Endotherms, as well as ectotherms inhabiting warm environments, have higher rates of deamination of 5mC nucleotides, resulting in lower global methylation levels and fewer CpG dinucleotides genome-wide (Sved and Bird 1990; Jabbari et al. 1997; Varriale and Bernardi 2006; Bernardi 2007; Bucciarelli et al. 2009). We use these data to test whether methylation levels and genome-wide CpG dinucleotide levels are driven by genome size and/or mode of body temperature regulation. In previous studies, endothermy was confounded with large genome size. The integration of the data from salamanders — which combine huge genome sizes with ectothermy — allows us to decouple genome size from metabolic heat production, revealing that genome size alone lacks explanatory power for methylation and CpG dinucleotide levels across vertebrates.

## Materials and Methods

### Sampling Design, Tissue Dissection, and DNA Extraction for Analysis of Global Methylation

Our first goal was to test for variation in methylation levels across tissues in salamanders. For this analysis, we focused on the model salamander *Ambystoma mexicanum*, widely used for studies of development and regeneration (Tank et al. 1976; Seifert et al. 2012; Keinath et al. 2015; Sessions and Wake 2020). Additionally, it is one of the few species of salamander to have its entire genome assembled and annotated (Keinath et al. 2015; Nowoshilow et al. 2018). We analyzed brain, heart, lung, liver, intestine, muscle, skin, and testes or ovaries from three adult males and three adult females.

Our second goal was to measure methylation levels in salamanders with gigantic genomes. For this analysis, we used muscle tissue as the single representative tissue, and we chose species that span the upper range of diploid genome sizes and ploidy in salamanders. The diploid species were *Plethodon cinereus* and *Necturus beyeri*, as well as *A. mexicanum. Plethodon cinereus* has a range of published genome sizes with an average of ~22 Gb (Gregory 2020), although our measurements indicate a genome size of 29.3 Gb (Itgen et al. 2021). The genome size of *Necturus beyeri* is unknown, but size estimates for congeners are 120 Gb (*N. lewisi* and *N. punctatus*) and ~85 Gb (*N. maculosus*) (Gregory 2020). *Ambystoma mexicanum* has a genome size of 32.15 Gb (Keinath et al. 2015). The polyploid taxa were triploid and tetraploid unisexual biotypes formed naturally via kleptogenesis (Bogart et al. 2007) including one haploid *A. laterale* genome (~29 Gb) and either two or three *A. jeffersonianum* genomes (haploid = ~29 Gb; ~87-116 Gb total). We also included the diploid *A. jeffersonianum*. Three individuals were sampled from each species/biotype (Table 1).

**Table 1.**
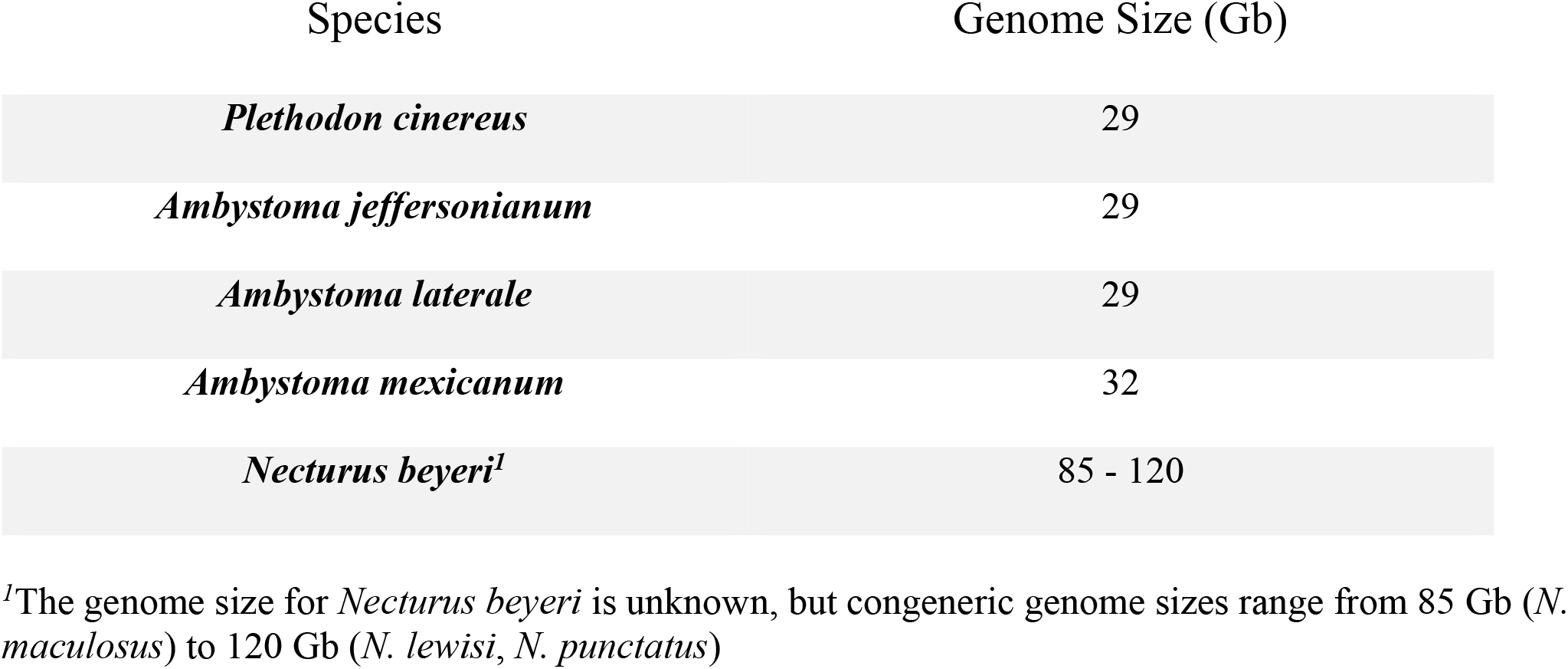
Genome sizes for salamanders sampled for global methylation

Our third goal was to compare methylation levels in salamanders with those from vertebrates with more typical genome sizes. For this analysis, we combined our new data with previously published methylation levels of brain and liver tissue from fish (*Carassius auratus*, *Perca flavescens*, *Salvelinus namaycush*), birds (*Coturnix japonica*, *Gallus gallus*), amphibians (*Xenopus laevis*, *Xenopus tropicalis*), reptiles (*Anolis carolinensis*), and mammals (*Delphinus delphis*, *Lagenorhynchus acutus*, *Neovison vison*, *Ursus maritimus*) (Basu et al. 2013; Head et al. 2014).

*Plethodon cinereus* were field collected between May and August of 2018 in South Cherry Valley and Oneonta, Otsego County, New York. *Necturus beyeri* were obtained commercially through a private vendor in November of 2019. *Ambystoma mexicanum* were obtained from the Ambystoma Genetic Stock Center in July of 2019. *Ambystoma jeffersonianum* and *Ambystoma laterale* were field collected from northern Kentucky (Kenton County) and Connecticut (Litchfield County). Polyploid taxa were collected from New Jersey (Morris County), Kentucky (Kenton County), and Ohio (Crawford County). All field collections took place in the spring of 2019 and 2020.

Specimens of *A. mexicanum*, *P. cinereus*, and *N. beyeri* were euthanized in MS222 and tissues were dissected and flash frozen in liquid nitrogen. DNA was extracted using a DNeasy Blood and Tissue extraction kit (Qiagen) following manufacturer instruction, including the optional RNase treatment step. DNA concentration and quality were measured using a Nanodrop spectrophotometer (Thermo Fisher Scientific). Tissue samples were stored at −80°C until use. For the polyploid *Ambystoma* samples, clippings of 1-3 mm of tissue from each animal’s tail were collected and stored immediately in 90% ethanol, then stored at −20°C until use. DNA was extracted using DNeasy Blood and Tissue extraction kits (Qiagen), using a double final elution step with a 25% decrease in volume to increase DNA concentration. DNA concentration was measured using a Qubit 3 fluorometer (ThermoFisher Scientific) with the dsDNA BR (Broad Range) assay. Ploidy and genomic composition of each sample were evaluated by sequencing mitochondrial loci and genotyping species-specific microsatellite markers as detailed in previous work (Denton et al. 2017). All work was carried out in accordance with either Colorado State University IACUC protocol number 17-7189A or University of Minnesota Morris IACUC protocol number 1901-36686A.

### Luminometric Methylation Assay and Data Standardization

To measure DNA methylation levels, we used the luminometric methylation assay (LUMA), a form of pyrosequencing that targets CpG dinucleotides capable of methylation (Karimi et al. 2006). LUMA runs were carried out by the sequencing facility EpigenDx (Hopkinton, MA, USA) using a PyroMark MD system from Qiagen. Duplicate LUMA runs were done for each DNA sample. All assays included four Lambda DNA standards with methylation percentages of 0, 50, 60, and 100% as internal controls. To account for any differences across assay runs, we calibrated against the internal controls using inverse regression calibration (Ott and Longnecker, 2015). Random effects were added per subject and per combination of subject and tissue type to limit batch effects and account for pseudoreplication of duplicate runs.

### Comparative Analyses of Global Methylation

First, we tested for differences in methylation levels among the *A. mexicanum* tissues using a mixed model ANOVA, with tissue as a fixed effect factor and individual and duplicate runs as random effect factors, followed by a Tukey HSD to test for significance between tissues. Next, we tested for differences in methylation levels among the polyploid unisexual salamander biotypes *A. laterale* (LJJ) and *A. laterale* (LJJJ) and the four diploid species *A. jeffersonianum* (JJ), *A. mexicanum*, *P. cinereus*, and *N. beyeri* using a mixed model ANOVA, with species as a fixed effect factor and individual and duplicate runs as random effect factors. We then performed a Tukey HSD to test for significance between species/biotype. Finally, we tested whether methylation levels vary across vertebrates as a function of salamander vs. non-salamander (i.e. genome size ≥ 29 Gb vs. genome size ≤ 6.4 Gb), ploidy (i.e. diploid vs. polyploid), and body temperature regulation (i.e. endothermy vs. ectothermy). We assigned species to the appropriate subgroup(s) and tested for variation between subgroups using linear regression contrasts. We carried out all analyses in R Studio (Martin 2021; Team 2021) using R packages emmeans (Lenth 2021), parameters (Lüdecke et al. 2020), and lme4 (Bates et al. 2015). We used the ggpubr package (Kassambara 2020) to visualize the results.

### Effects of Global Methylation and Genome Size on CpG Dinucleotide Levels

Our final goal was to test whether enormous genome sizes translate into changes in the nucleotide compositional landscape, as predicted based on the relationships among genome size, transposable element load, methylation level, and C ➔ T deamination mutation inferred across 53 animals (47 vertebrates, 6 invertebrates) with more typical genome sizes (89 Mb to 4 Gb) (Zhou et al. 2020). C ➔ T deamination mutations occur preferentially at methylated CpG dinucleotides (Ehrlich and Wang 1981); thus, higher levels of methylation, associated with methylation-based silencing of a greater TE load, are predicted to correlate with fewer CpG dinucleotides genome-wide than are expected based on nucleotide frequencies (Zhou et al. 2020). We obtained publicly available genomic sequence data for nine species of salamanders with varying genome sizes: *Desmognathus ochrophaeus* (~15 Gb; SRA046120.1), *Pleurodeles waltl* (~25 Gb; DRX131369), *Eurycea tynerensis* (~25 Gb; SRA046121.1), *Batrachoseps nigriventris* (~26 Gb; SRA046116.1), *Ambystoma mexicanum* (~30 Gb; ASM291563v2), *Bolitoglossa occidentalis* (~43.5 Gb; SRA046118.1)*, Aneides flavipunctatus* (~45 Gb; SRA046114.1)*, Bolitoglossa rostrata* (~48 Gb; SRA046119.1), and *Cryptobranchus alleganiesis* (~55 Gb; SRA073787) (Gregory 2020). *Ambystoma mexicanum* has a well-assembled genome (including the repetitive regions) based on deep, short-read coverage, long reads, and optical mapping (Nowoshilow et al. 2018). *Pleurodeles waltl* has a short-read Illumina assembly, with the repeat elements constructed through a majority vote k-mer extension algorithim (Elewa et al. 2017); however, we used the unassembled Illumina HiSeq 2000 trimmed and quality filtered reads (~0.25X coverage) to avoid bias introduced by the relative ease of assembling genic versus repetitive sequences. Datasets for the remaining species are all low-coverage 454 shotgun reads representing about 0.1% - 1% of the total genome (Sun et al. 2012; Sun and Mueller 2014). Each dataset was run through a pipeline bash script that removed tags and counted the numbers of each individual nucleotide and CpG dinucleotides present. The observed/expected ratio (O/E) of CpG dinucleotides was calculated as the observed number of CpG dinucleotides, CpG/N, divided by the expected number of CpG dinucleotides, (C × G)/(N × N). C is observed cytosines, G is observed guanines, CpG is the observed CpG dinucleotides, and N is the total number of base pairs. Overall CpG O/E = (CpG/N)/((CxG)/(N^2^)) (Zhou et al. 2020).

We tested for a relationship between genome size and CpG O/E dinucleotides among the nine salamander species using a linear regression analysis. We also corrected for phylogeny using phylogenetic independent contrasts (PIC) with the tree for the nine focal species subsampled from a comprehensive amphibian phylogeny (Pyron and Wiens, 2011). PIC analyses were carried out using R packages ape (Paradis and Schliep 2019), Geiger (Harmon et al. 2008), nlme (Pinheiro et al. 2020), and phylools (Revell 2012). Next, we compared the salamander CpG O/E dinucleotide values to all species published in (Zhou et al. 2020); this study included full genome assemblies from species with genome sizes ranging from 0.36 to 4.78 Gb and showed a negative correlation between genome size and CpG O/E (Zhou et al. 2020). To ensure that the datasets were comparable, we subsampled Illumina reads of 20 species from the Zhou et al. dataset to generate datasets with coverage that approximated the coverage of the salamander datasets, and we calculated CpG O/E from these subsampled datasets. We then compared the CpG O/E values based on full genome assemblies to those we calculated from subsampled short read datasets using linear regression and found a significant positive correlation between the two sets of estimates (p < 0.00001, R = 0.83; subsampled estimates equally likely to be slightly above or below whole-geome estimates), suggesting that comparisons across these different datasets are unlikely to introduce bias due to comparing whole-genome assemblies to unassembled read datasets.

## Results

### Methylation Levels are Lower in the Lungs

Methylation levels significantly differ between some tissues: Brain – Lungs (p = 0.003), Gill – Lungs (p = 0.012), Heart – Lungs (p = 0.012), and Liver – Lungs (p = 0.001) (df = 8 and an overall F-value of 4.5) (Figure 1). Lungs were not used for downstream across-species comparisons.

**Fig. 1.**
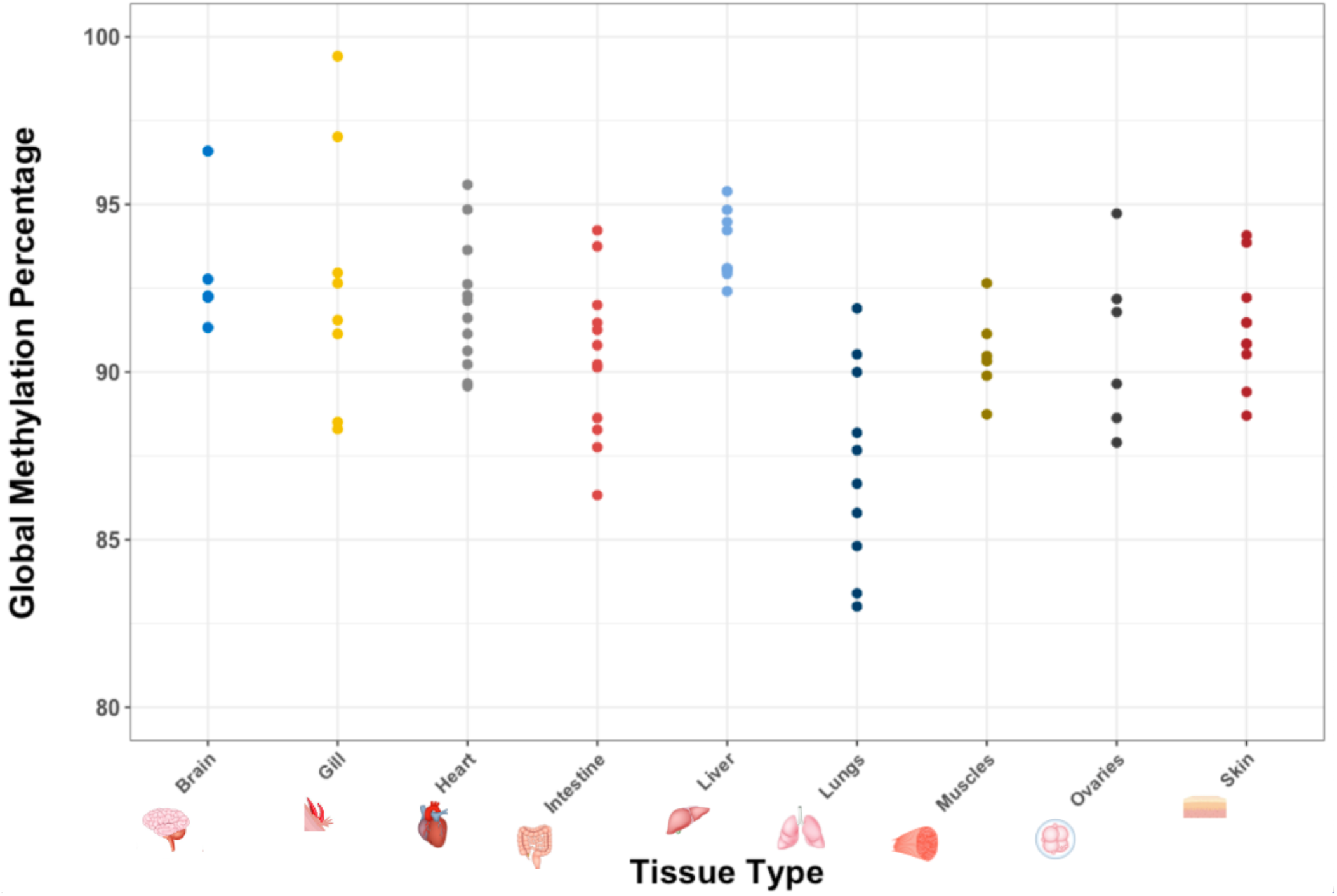
Methylation levels (percentage methylated cytosines at CpG dinucleotide sites) of DNA extracted from different tissues in *Ambystoma mexicanum*. Lungs have significantly lower methylation levels compared to some tissues: Brain – Lungs, Heart – Lungs, Gill – Lungs, and Liver – Lungs.

### Methylation Levels Do Not Vary Across Salamander Species

Methylation levels are not significantly different between salamander species, despite differences in diploid genome size and ploidy (df = 5, F-value = 1.11, p = 0.4; Figure 2). We note that the two species with the most nuclear DNA — the tetraploid *Ambystoma* biotype LJJJ and the diploid *Necturus beyeri —* have the largest intraspecific variances.

**Fig. 2.**
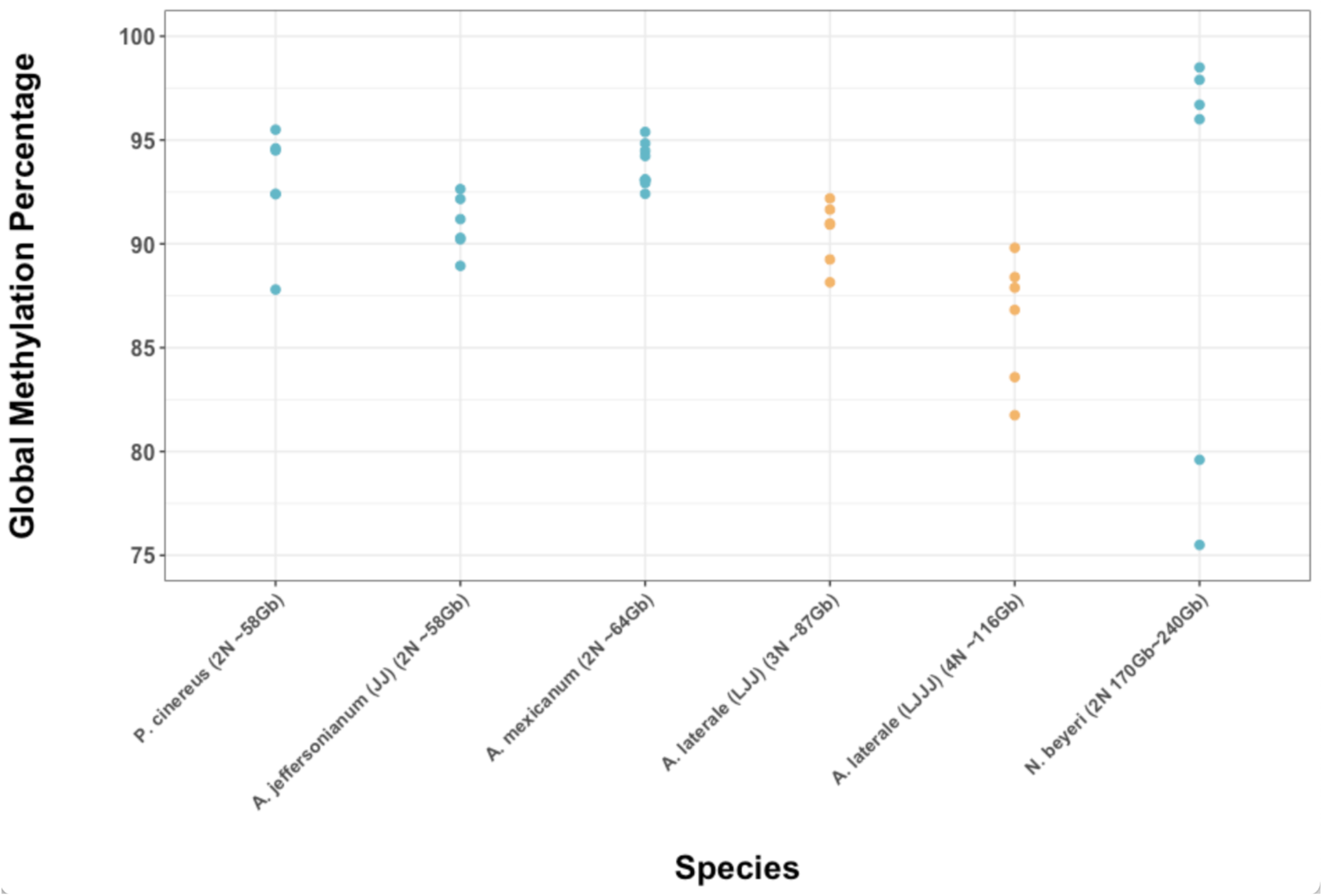
Methylation levels (percentage of methylated cytosines at CpG dinucleotide sites) of six different species/biotypes of salamanders with different amounts of nuclear DNA. Teal represents diploid species and orange represents polyploid species. There are no significant differences among groups.

### Across Vertebrates, Methylation Levels Reflect Body Temperature Regulation, Not Nuclear DNA Content

Ectotherms as a whole (including salamanders) have significantly higher levels of methylation than endotherms (df = 108, t-ratio = −15.84, p < 0.001, Figure 3). Salamanders themselves have methylation levels that are not significantly different from other ectotherms (df = 108, t-ratio = 0.88, p = 0.38), despite their enormous genome sizes. Excluding salamanders, there is still a significant difference between ectotherms and endotherms (df = 108, t-ratio = − 9.47, p < 0.001). Polyploidy (seen in *X. laevis* and the two uniexual *Ambystoma* biotypes) does not significantly affect methylation levels (df = 108, t-ratio = 0.38, p = 0.71), although tetraploid *X. laevis* has higher methylation than diploid *X. tropicalis*. This may reflect the fact that *X. laevis* is a 30-million-year-old polyploidization event (Tymowska and Fischberg 1973; Hughes and Hughes 1993), whereas polyploidization happens anew each generation in *Ambystoma*.

**Fig. 3.**
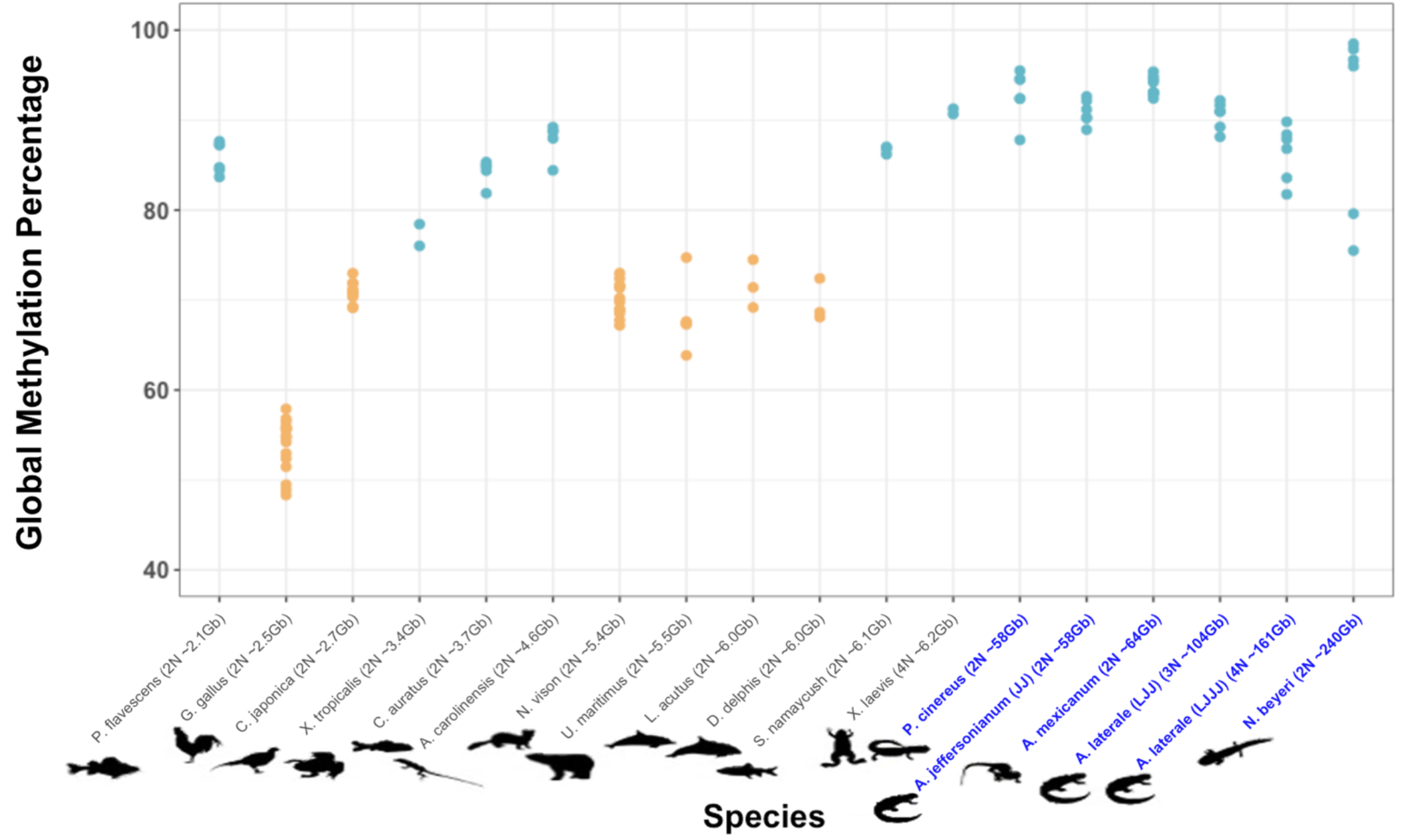
Methylation levels (percentage of methylated cytosines at CpG dinucleotide sites) across vertebrates. Diploid, triploid, or tetraploid genome sizes are listed with species names. Orange dots indicate endothermy and teal dots indicate ectothermy. Blue font indicates salamanders. Ectotherms have higher methylation levels than endotherms (p < 0.001).

### Large, Highly Methylated Genomes Show No Relationship Between Deamination-Driven Transition Mutations and Genome Size

The observed versus expected ratios of CpG dinucleotides (CpG O/E) for nine species of salamanders range from 0.47 – 1.08. There is no significant relationship between genome size and CpG O/E dinucleotides across salamanders, despite a ~3-fold difference in genome size (Figure 4). We note that *C. alleganiensis* is an a statistical outlier, with an O/E CpG value of 1.08 that invites further study; however, our conclusions are not affected by this data point. Overall, the range of CpG O/E values in salamanders is higher than those seen in the largest genomes sampled in previous analyses of tetrapods (2 Gb – 4 Gb, CpG O/E range 0.13 – 0.54) (Zhou et al. 2020), suggesting that the negative correlation between genome size and CpG O/E demonstrated across a smaller range of animal genome sizes does not hold across the full range of animal genome size.

**Fig. 4.**
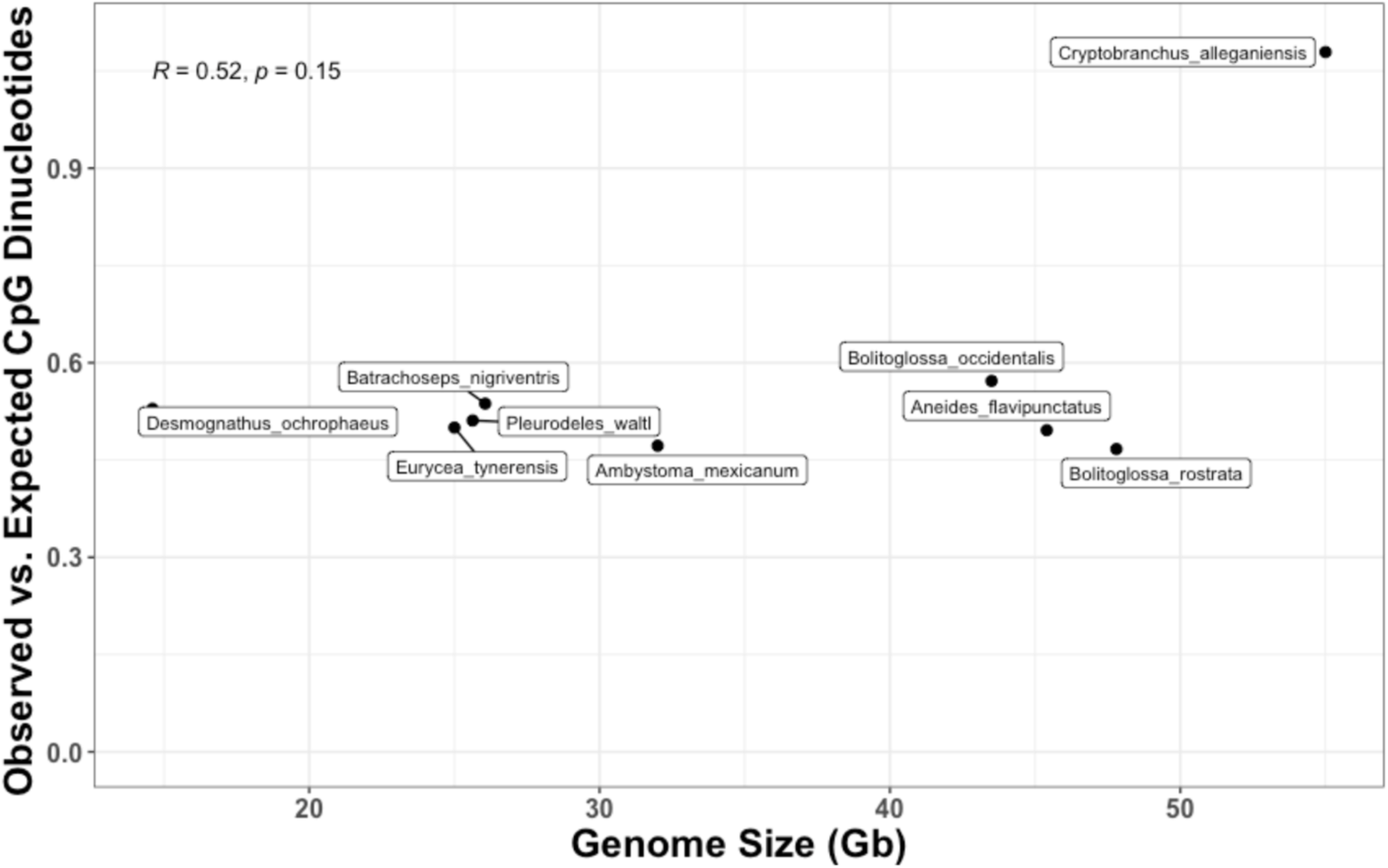
The observed versus expected ratios of CpG dinucleotides (O/E) for nine species of salamanders. There is no significant correlation between genome size and O/E CpG (p = 0.15)

## Discussion

### Body Temperature, Not Nuclear DNA Content, is the Main Predictor of Methylation Levels and CpG Dinucleotide Levels Across Vertebrate Genomes

Despite spanning a ~100-fold difference in genome size, the methylation levels of ectotherms were not significantly different from one another. Additionally, the methylation levels of polyploids were not significantly different from diploids. In contrast, endotherms had significantly lower levels of methylation compared to ectotherms, with or without the inclusion of salamanders. These results suggest that the largest predictor of DNA methylation levels in vertebrates is the maintenance of a high body temperature by the production of metabolic heat, and not the amount of DNA in the nuclear genome.

Endothermy and ectothermy have been proposed as the primary determinants of global methylation levels in vertebrates before (Jabbari et al. 1997). Mammals and birds, which have high, endothermically-maintained body temperatures, have lower methylation levels than amphibians and reptiles, which have lower body temperatures. This difference in methylation is paralleled in congeneric species of gobies that live in vastly different temperatures in the Gulf of California—the species that lives in warmer temperatures has lower levels of DNA methylation (Bucciarelli et al. 2009). Similarly, temperate and tropical fish have lower methylation levels than polar fish (Varriale and Bernardi 2006). In all of these cases, the lower methylation levels found in animals with higher body temperatures likely reflect the higher rates of 5mC deamination of CpG dinucleotides at warmer temperatures, leading to faster loss of methylated cytosines from the genome (Jabbari et al. 1997; Varriale and Bernardi 2006; Bernardi 2007; Bucciarelli et al. 2009). High methylation levels likely evolved at the base of vertebrates and remain high in ectotherms, irrespective of genome size; the independent evolutionary acquisitions of endothermy resulted in lower methylation levels.

5mC deamination rates should also impact the global nucleotide composition landscape because deamination causes transition mutations from C to T. Thus, endothermic vertebrates, as well as ectotherms living at high temperatures, should have fewer CpG dinucleotide sites than expected based on their nucleotide frequencies (lower O/E CpG values). Our results, in combination with those of Zhou et al. (2020), suggest that endothermy and ectothermy in vertebrates are also the primary predictors of CpG O/E values. Our salamander CpG O/E values broadly overlap with those of the 14 fish, 3 reptile, and 1 frog species sampled by Zhou et al, with genome sizes ranging from 0.36 – 2.86 Gb. In contrast, the 5 bird and 88 mammal species sampled by Zhou et al., with genome sizes ranging from 1.06 – 4.78 Gb, have lower CpG O/E, as predicted by their endothermic body temperature regulation. We re-analyzed the data from Zhou et al, running a linear regression with first genome size as the predictor, followed by ectothermy/endothermy. The R^2^ value increased from 0.26 (when using genome size as the predictor) to 0.48 (when using ectothermy vs. endothermy). This suggests that much of the signal within the Zhou et al. dataset comes from the fact that mammals are endothermic and also tend to have larger genomes than fish, reptiles, and frogs—but it is endothermy itself, rather than genome size, that is driving the CpG O/E pattern (Supplemental Figures 1 and 2). Within the non-salamander ectotherms alone, however, genome size is negatively correlated with CpG O/E based on linear regression analysis (p = 0.009), suggesting a possible relationship between TE load, methylation, and CpG loss in ectotherms with relatively small genomes that warrants further investigation.

Given these results, is there a potential mechanistic relationship between endothermy and the slightly increased TE load of mammals? Higher deamination rates would decrease methylation of TE sequences in mammals, which might negatively impact their silencing and allow for greater activity. In addition, higher deamination rates would yield higher transition mutation rates of TE sequences, which in turn could have two outcomes: 1) the production of divergent TE sequences that escape sequence-specific TE silencing, leading to greater TE activity, and/or 2) the production of TE sequences that lose their functional ORFs, rendering them incapable of autonomous transposition and leading to decreased TE activity. There are also groups that do not show an association between endothermy and increased TE load and genome size. Birds — also endothermic — have lower TE loads and smaller genome sizes, and salamanders — ectotherms — have the highest TE loads and largest genomes among tetrapods. In total, TE activity and abundance reflect the interaction of diverse forces, which may or may not include a relationship between methylation, mutation, and TE silencing that contributes to mammals’ slightly larger genome sizes relative to non-salamander vertebrates.

## Acknowledgments

We thank M Itgen for *P. cinereus* tissues. We thank W Zhou, G Liang, PL Molloyd, and PA Jones for providing their full dataset for analysis. We thank members of A Adams’ dissertation committee K Hoke, D Sloan, and J Hansen for helpful discussion and feedback throughout the project. We thank the *Ambystoma* Genetic Stock Center, funded by NIH Grant P40-OD019794. Funding for this project was provided by NSF grant 1911585 to R Mueller, a UMN Grant-in-Aid award and NSF grant 2045704 to R Denton, and by Colorado State University.

---

**Supplementary Fig. 1.**
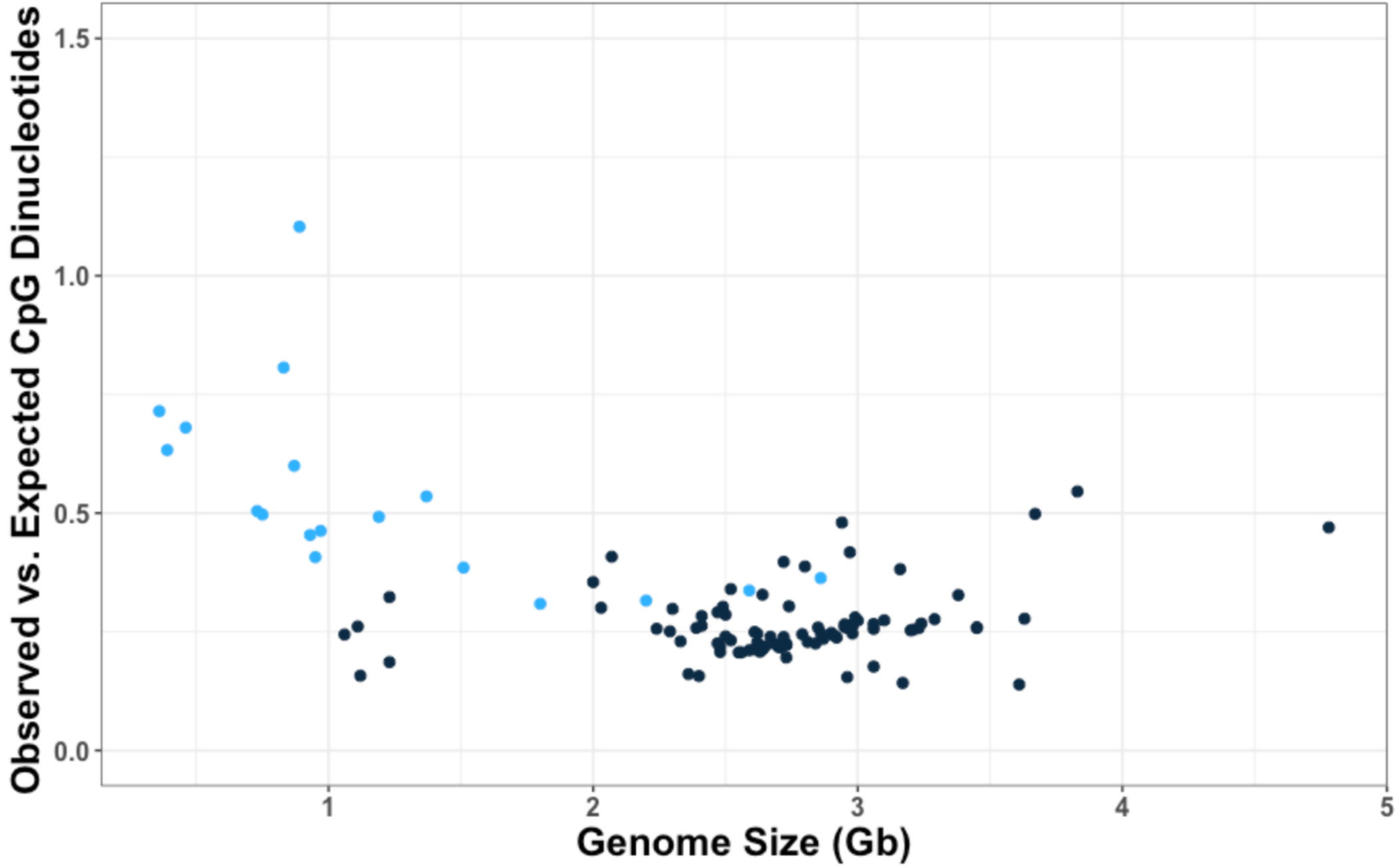
Data from Zhou et al. 2020. Each dot on the graph represents a vertebrate species. Blue dots are ectotherms and black dots are endotherms. Regression analysis shows a significant negative correlation (p = 0.009) between genome size and O/E for the ectotherms. There is no significant relationship between genome size and O/E for endotherms.

**Supplementary Fig. 2.**
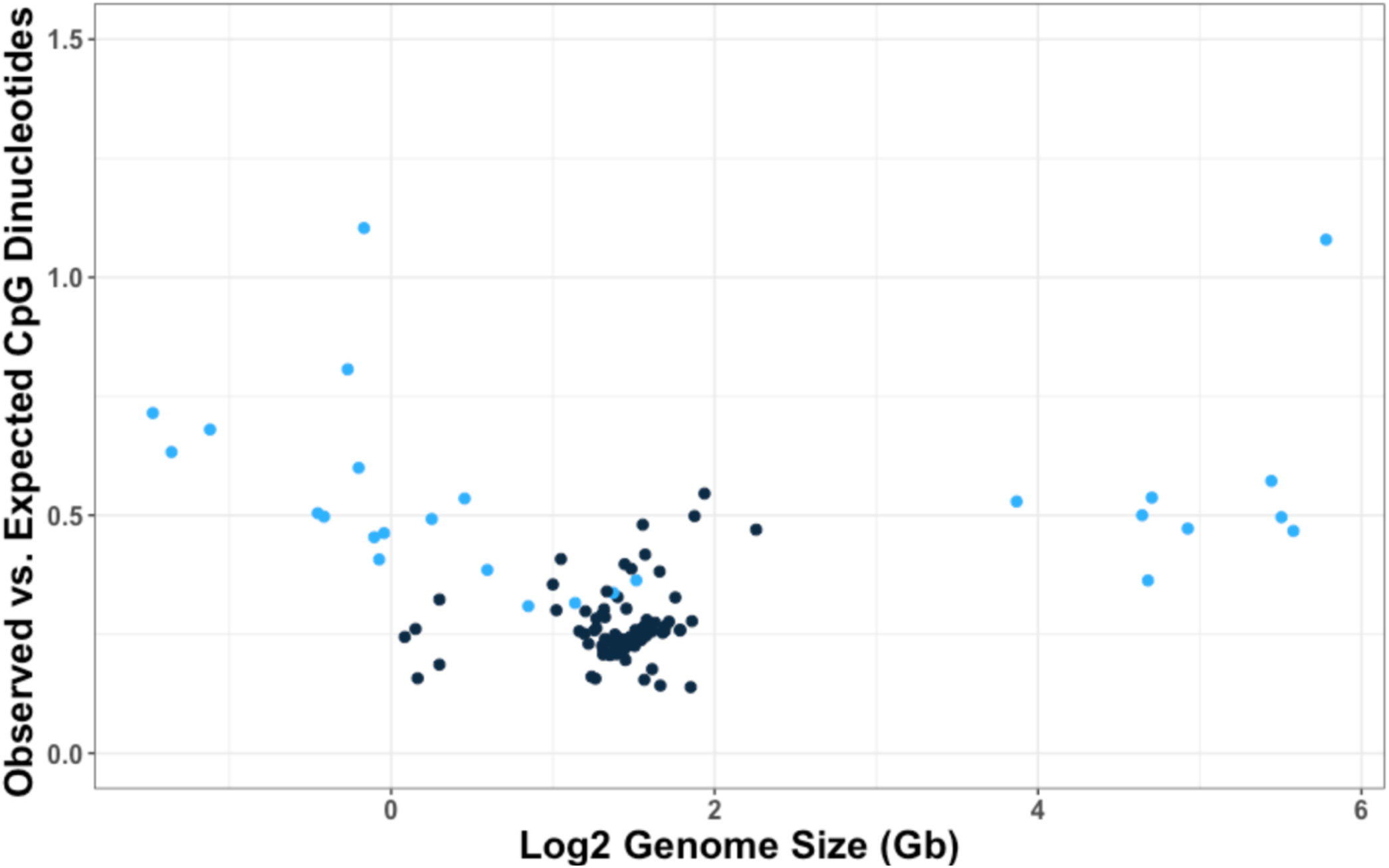
Data from Zhou et al. 2020 and data from 9 different species of salamander from this study. Genome size is shown on a log scale. Blue dots are ectotherms (non-salamander log2 genome size < 2, salamander log2 genome size > 3) and black dots are endotherms. Regression analysis shows no significant correlation (p = 0.4) between genome size and O/E for the ectotherms including the salamanders.

